# Measurements of three-dimensional refractive index tomography and membrane deformability of live erythrocytes from *Pelophylax nigromaculatus*

**DOI:** 10.1101/206821

**Authors:** Geon Kim, Moosung Lee, SeongYeon Youn, EuiTae Lee, Daeheon Kwon, Jonghun Shin, SangYun Lee, Youn Sil Lee, YongKeun Park

## Abstract

Unlike mammalian erythrocytes, amphibian erythrocytes have distinct morphological features including large cell sizes and the presence of nuclei. The sizes of the cytoplasm and nuclei of erythrocytes vary significantly over different species, their environments, or pathophysiology, which makes hematological studies important for investigating amphibian species. Here, we present a label-free three-dimensional optical quantification of individual amphibian erythrocytes from frogs *Pelophylax nigromaculatus (Rana nigromaculata)*. Using optical diffraction tomography, we measured three-dimensional refractive index (RI) tomograms of the cells, which clearly distinguished the cytoplasm and nuclei inside the erythrocytes. From the measured RI tomograms, we extracted the relevant biochemical parameters of the cells, including hemoglobin contents and hemoglobin concentrations. Furthermore, we measured dynamic membrane fluctuations and investigated the mechanical properties of the cell membrane. From the statistical and correlative analysis of these retrieved parameters, we investigated interspecific differences between frogs and previously studied mammals.

## Introduction

Hematological studies of amphibians play an important role in the study of their evolutionary processes following the vertebrate invasion of the land^1^. One of their critical traits of amphibians, compared to other vertebrates, is their unusually large erythrocytes, which may be associated with the adaptation for the transition from aquatic to terrestrial life. Because of their distinct morphologies, amphibian erythrocytes have been extensively investigated. Previous studies have suggested that the adaptive structures of the cells are dependent on both intrinsic factors including ages and species^2,3^, and extrinsic factors including habitat conditions^4^.

To quantitatively assess the various physiological properties of the cells, various experimental methods have been exploited. For example, bright-field microscopy has been used to quantify the size parameters of the cells in two-dimensions (2D), such as their aspect ratios^5^. However, this method has limitations when assessing the three-dimensional (3D) information of a cell. In addition, the method often relies on labeling methods such as fluorescent proteins or organic dyes, which may perturb the physiological conditions of the cells. Alternatively, a complete blood count was used for a high-throughput measurement of both morphological and biochemical parameters^6^. Electron microscopy was also implemented to visualize viral particles inside erythrocytes beyond the optical diffraction limit^7^. However, these methods present challenges when studying the dynamics of live individual cells *in vitro*.

Recently, quantitative phase imaging (QPI) techniques have been developed as a label-free imaging method for the study of live cells and tissues^8,9^. QPI employs the principle of laser interferometric microscopy or digital holographic microscopy to measure the optical phase delay induced by a transparent specimen. Because the optical phase delay of a biological cell is linearly proportional to its thickness and relative refractive index (RI)^10^, QPI provides a label-free, quantitative imaging capability with nanometer sensitivity. In particular, the morphology and biophysical properties of human erythrocytes and their pathophysiology have been extensively studied using this method^11–15^.

In order to reconstruct 3D tomograms of unlabeled transparent cells, considerable advances have been made in tomographic QPI techniques^16–20^. Among them, optical diffraction tomography (ODT) or holotomography enables to reconstruct the 3D RI tomogram of a sample from multiple 2D QPI images with various illumination angles^18,21–23^. ODT has shown potential for precisely assessing the 3D morphological properties of blood cells at the individual cell level, and their related pathophysiologies, such as malaria or babesia infection^18,24–27^. However, although ODT has been previously used to study mammalian blood cells^28–32^, its potential for studying amphibian RBCs has not been investigated.

Here, we present 3D optical tomographic measurements of individual amphibian erythrocytes from frogs *Pelophylax nigromaculatus (Rana nigromaculata).* Using ODT, 3D RI tomograms of erythrocytes were measured at the individual cell level, which clearly visualized the 3D structures of the cytoplasm and nuclei, and provided morphological information such as cell volume and surface area, nucleus volume, surface area, and sphericity index. Exploiting the measured RI values, biochemical properties of the cells including hemoglobin (Hb) concentrations and contents were also retrieved. Furthermore, dynamic membrane fluctuations of individual live cells were measured, providing data on the biomechanical properties of the cell membranes. From the measured cellular parameters, we were able to compare the interspecific differences of erythrocytes from amphibians and formerly investigated mammals. Also, we examined the correlative relations between the retrieved parameters, which suggested the general characteristics of erythrocytes from different species.

## Results and Discussions

### 3D RI tomograms of frog erythrocytes

The principle of ODT is illustrated in Fig. 1a. Similar to X-ray computed tomography (CT), ODT reconstructs the 3D tomogram of a sample from multiple 2D images of the sample obtained with various illumination angles^22^. While CT measures X-ray absorptivity in 3D, ODT measures RI tomography containing information about both the sample-induced light absorption and refraction.

**Figure 1.**
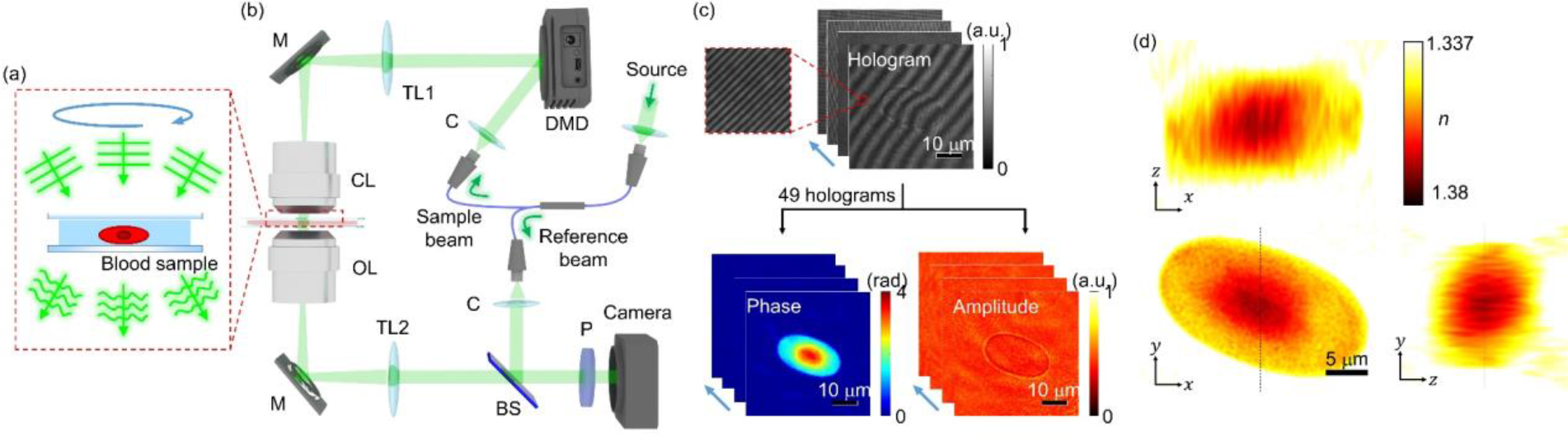
Principles of reconstructing 3D RI tomograms of frog erythrocytes using ODT. **(a)** Schematic diagram of image acquisition process in ODT. **(b)** Experimental setup. C: collimator; DMD: digital micromirror device; TL: tube lens; M: mirror, CL: condenser lens; OL: objective lens, BS: beam splitter; P: polarizer. **(c)** 49 measured off-axis holograms and retrieved phase and amplitude maps. **(d)** Cross sections of the reconstructed RI tomogram from the phase and amplitude maps in (c).

In order to measure the 3D RI tomograms of frog erythrocytes, we utilized a commercial ODT setup^33,34^ (HT-1S and HT-2S, Tomocube Inc.) [Fig. 1b]. The ODT setup is based on an off-axis Mach-Zehnder interferometer equipped with a digital micromirror device (DMD). A 2×2 single-mode fiber coupler splits a coherent, monochromatic laser (wavelength, *λ* = 532 nm) into a sample and a reference arm, respectively. To control the illumination angle of the beam impinging onto a sample, a DMD is utilized to diffract light into various angles^35,36^. The light scattered by the sample is then transmitted through an objective lens (60×, numerical aperture = 0.9) and a tube lens (*f* = 175 mm). The sample beam is combined with the reference beam by a beam splitter and filtered by a linear polarizer. The resultant spatially modulated hologram is recorded by an image sensor. The image sensor is synchronized with the DMD to record 49 holograms of the sample illuminated with different angles (Fig. 1c). Using a phase retrieval algorithm^37,38^, the amplitude and phase images are retrieved from the measured holograms. Based on the Fourier diffraction theorem with Rytov approximation^39^, the 3D RI tomogram of the sample is reconstructed from the retrieved amplitude and phase images (Fig. 1d). The lateral and axial optical resolutions of the ODT system were 166 nm and 1 μm respectively, according to the Lauer criterion^40^. Detailed information about the system and reconstruction algorithms can also be found elsewhere^18,33,41^.

### Anatomic structures of frog erythrocytes

Figure 2 presents the representative 3D RI structure of a frog erythrocyte. The cross-sections of the RI tomogram clearly show the ellipsoidal shape of a frog erythrocyte (Fig. 2a), which significantly differs from the biconcave shapes of mammalian erythrocytes. Also, the frog erythrocyte presents a central region whose RI values were higher than the outer region. The bright-field and 3D fluorescence images of the same sample stained with DAPI clearly indicate that this central area corresponds to a nucleus (Fig. 2b).

**Figure 2.**
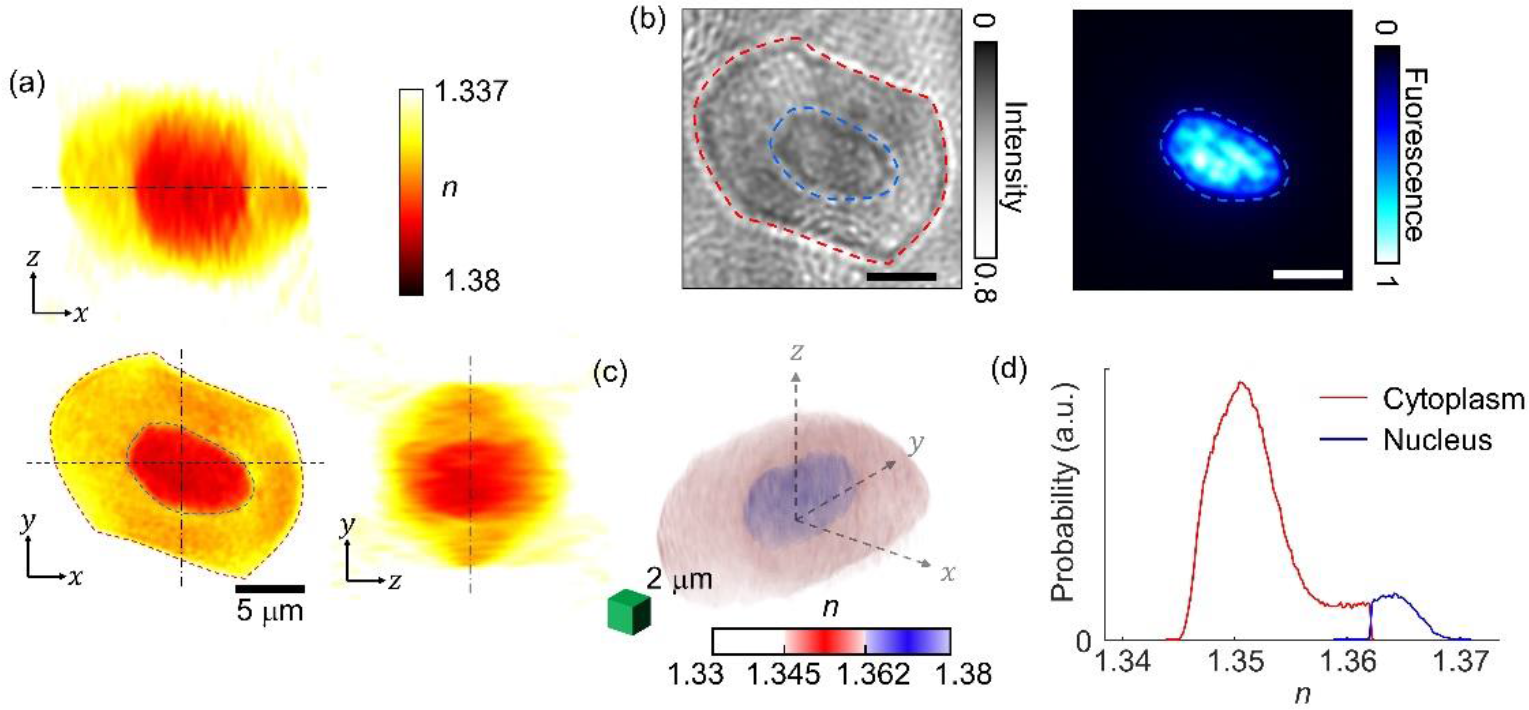
Structural analyses of frog erythrocytes using 3D RI maps. **(a)** Cross sections of the RI map of a frog erythrocyte. The red-and blue-dashed regions correspond to the cytoplasm and nucleus, respectively. **(b)** Bright-field image and DAPI fluorescence image for cross-validation of the cytoplasm and nucleus. **(c)** 3D-rendered isosurface image of the RI tomogram. **(d)** RI histograms for the cytoplasm and nucleus in (c).

For quantitative analysis of the nucleus, the RI threshold between the cytoplasm and nucleus was determined to be 1.362, and was selected by comparing both the focal planes of the 3D RI tomograms and the 3D fluorescence images of the DAPI-stained cells. The cross-sectional images of the two images in different modalities showed a 93% correlation, verifying the RI threshold. Using the RI criterion, the cytoplasm and the nucleus of individual frog erythrocytes were segmented in 3D (Fig. 2c). The 3D isosurface image clearly visualizes the nucleus located in the middle of the cell. In addition, the averaged RI values of the nucleus and cytoplasm of the erythrocyte in Figs. 2a-c were measured and determined to be 1.365 ± 0.002 and 1.351 ± 0.004, respectively (Fig. 2d).

### Morphological properties of frog erythrocytes

For a statistical analysis of the quantitative parameters of individual frog erythrocytes, we retrieved the volumes, surface areas, SIs, Hb contents, and Hb concentrations of 128 erythrocytes from six frogs (Fig. 3). For interspecies analyses, we also compared each parameter of the frog erythrocytes with those of human and mouse erythrocytes, extracted from previous reports^26,42^.

**Figure 3.**
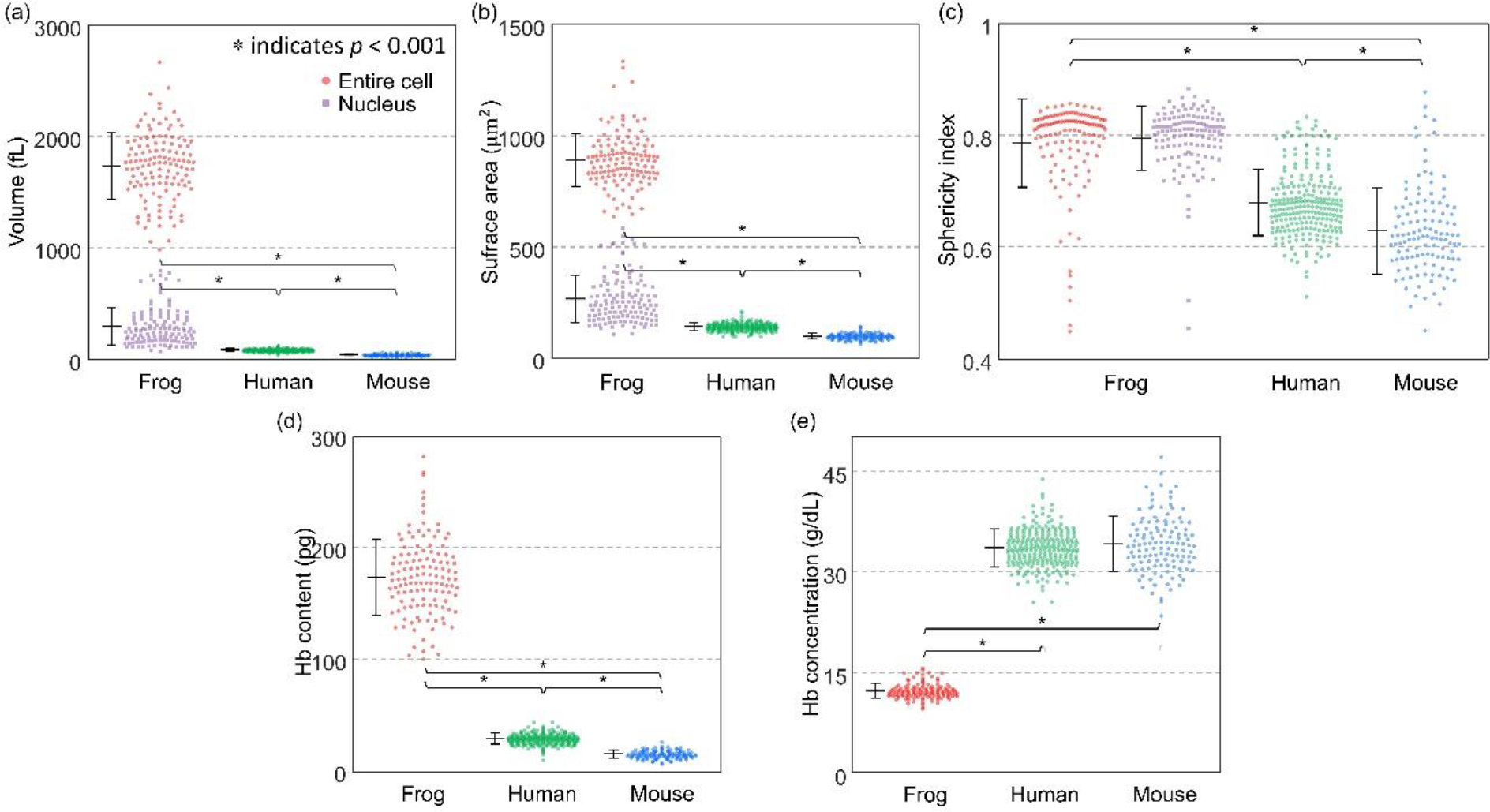
Statistical analyses of erythrocyte morphological and biochemical parameters for frogs, humans, and mice. **(a-c)** Distributions of morphological parameters including **(a)** volumes, **(b)** surface areas, and **(c)** sphericity indices. **(d-e)** Distributions of biochemical parameters including **(d)** Hb contents and **(e)** Hb concentrations. * indicates p < 0.001.

As is well known in 2D microscopy^3^, frog erythrocytes exhibit distinct morphological parameters that are unlike human or mouse erythrocytes. The mean values of the cellular volumes of the frog erythrocytes were 1737.46 ± 299.32 fL, which was an order of magnitude larger than that of human (90.5 ± 11.4 fL) and of mouse erythrocytes (45.6 ± 6.3 fL). Nucleus volumes of the amphibian cells were also measured to be 300.15 ± 167.07 fL, accounting for approximately 17% of the total erythrocyte volume. The mean values of the cellular surface area of the frog erythrocytes were 892.35 ± 119.07 μm^2^, which was about six to nine times larger than human (144.10 ± 17.40 μm^2^) and mouse erythrocytes (102.00 ± 13.00 μm^2^). The nucleus surface areas were on average 267.88 ± 106.39 μm^2^, approximately 30% smaller than the cellular area.

To investigate size independent morphological properties of the erythrocytes, we measured their sphericity indexes, the unitless parameters which quantify the shape resemblance to a sphere (Fig. 3c; see also *Methods*). The sphericity index is closer to one when the shape is closer to a sphere. The nuclei in the frog erythrocytes resembled ellipsoids; the average sphericity index of the nuclei was 0.80 ± 0.06. Like the nuclei, the membrane structures of the frog erythrocytes exhibited ellipsoidal shapes, whereas human and mouse erythrocytes exhibit biconcave shapes. The average values of sphericity indexes were calculated to be 0.79 ± 0.08, 0.68 ± 0.06, and 0.63 ± 0.08 for the frog, human, and mouse erythrocytes, respectively.

### Biochemical properties of frog erythrocytes

The biochemical properties of the cytoplasm, Hb concentrations, and contents were also obtained from the measured RI values (Figure 3d; see also Methods). The result showed that the frog erythrocytes had significantly higher amounts of Hb contents than human and mouse erythrocytes. The retrieved Hb contents were 174.35 ± 33.93, 30.3 ± 4.8, and 16.40 ± 3.50 pg for frogs, humans, and mice, respectively. The Hb contents of the frog erythrocytes measured by RI values were consistent with the previously estimated values for *Rana* frogs using the Sahli method^43^. The results showed that Hb concentrations in the frog groups were approximately three times lower than those of human and mouse groups (Fig. 3e). The measured Hb concentrations were 12.25 ± 1.12, 33.4 ± 2.8 and 34.0 ± 4.0 g/dL for frogs, humans, and mice, respectively. Note that these estimated low Hb concentrations of frog erythrocytes agree well with the previously reported values for mature *Rana* frogs^44^.

### Membrane deformability of frog erythrocytes

In order to investigate the membrane properties of the frog erythrocytes, we measured and quantified the dynamic fluctuations in the cell membrane (Fig. 4, see *Methods*). Membrane fluctuations of erythrocytes are strongly correlated with viscoelastic properties of the lipid membrane, spectrin network, and cytoplasmic viscosity, which can also be altered by diseases^11,12,14,42,45^.

**Figure 4.**
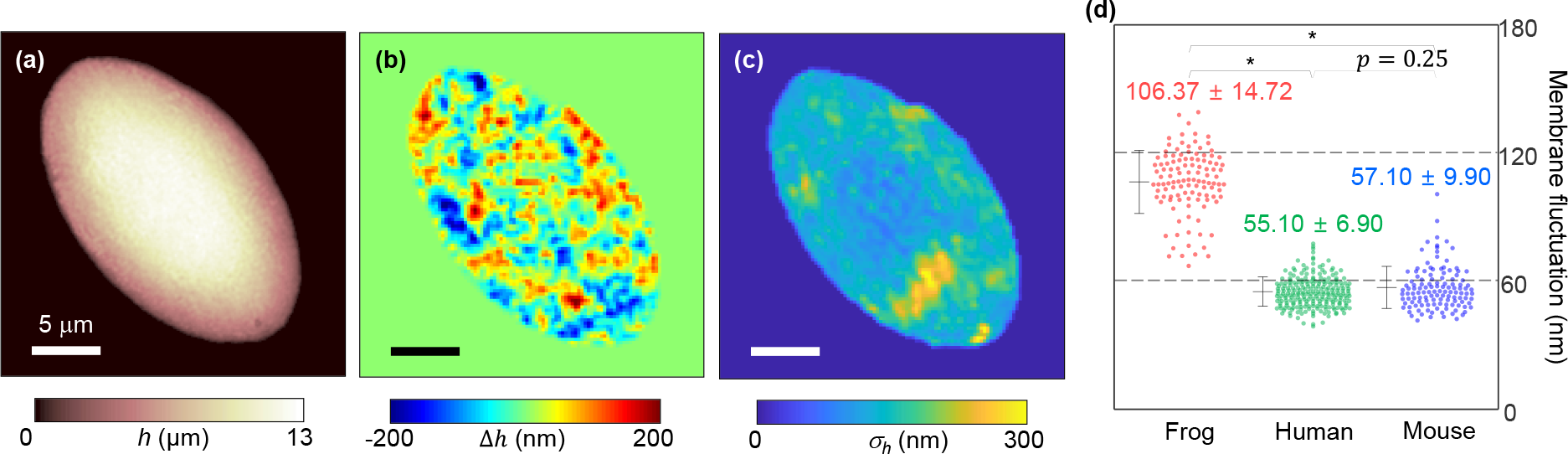
Measurement of temporal membrane fluctuation. **(a)** The thickness profile of a frog erythrocyte. **(b)** Instantaneous thickness shift of (a). **(c)** Membrane fluctuation map of (a). **(d)** Distributions of temporal membrane fluctuations of frogs, humans, and mice. * indicates p < 0.001.

Figures 4a and 4b show the height and the height change in a frog erythrocyte, respectively. The relative height change for 8 ms shows the dynamic membrane fluctuations of the sample (Fig. 4b). We defined the spatially resolved membrane fluctuation profiles as the standard deviations of the temporal height distributions, as in Fig. 4c. The membrane fluctuation map shows an inhomogeneous amplitude distribution over the cell membrane; the amplitude of the membrane fluctuation is slightly decreased near the nucleus. Considering cell structures, the decreased fluctuation near the nucleus can be explained by the presence of a stiff nucleus or cytoskeleton network anchoring the nucleus, or both^46^. For further investigation, correlative imaging with fluorescence microscopy would be helpful^34,47^.

Then we compared the membrane fluctuations of frog, human, and mouse erythrocytes (Fig. 4d). The results showed that the membrane fluctuation of frog erythrocytes was approximately twice greater than those of human and mouse erythrocytes. The dynamics membrane fluctuations σ_*h*_ were 106.37 ± 14.72 nm, 55.10 ± 6.90 nm, and 57.10 ± 9.90 nm for frogs, humans, and mice, respectively.

### Correlative analysis of quantitative parameters of frog erythrocytes

We further examined the physiological characteristics of frog erythrocytes via correlative analyses of the retrieved parameters (Fig. 5). The scatterplot between the cell volumes and the Hb contents show a positive linear correlation (Fig. 5a). The slopes obtained using least square fitting clearly indicate that for each species, Hb concentrations in individual cells do not vary significantly, regardless of the cell size. Although Hb concentrations in the human and mouse groups showed similar values (33.5 and 34.0 g/dL, respectively), the Hb concentration of the frog group showed a significantly lower value (10.0 g/dL). This result implies that homeostasis mechanisms for Hb concentration in amphibian erythrocytes differ from those of mammalian cells^48,49^.

**Figure 5.**
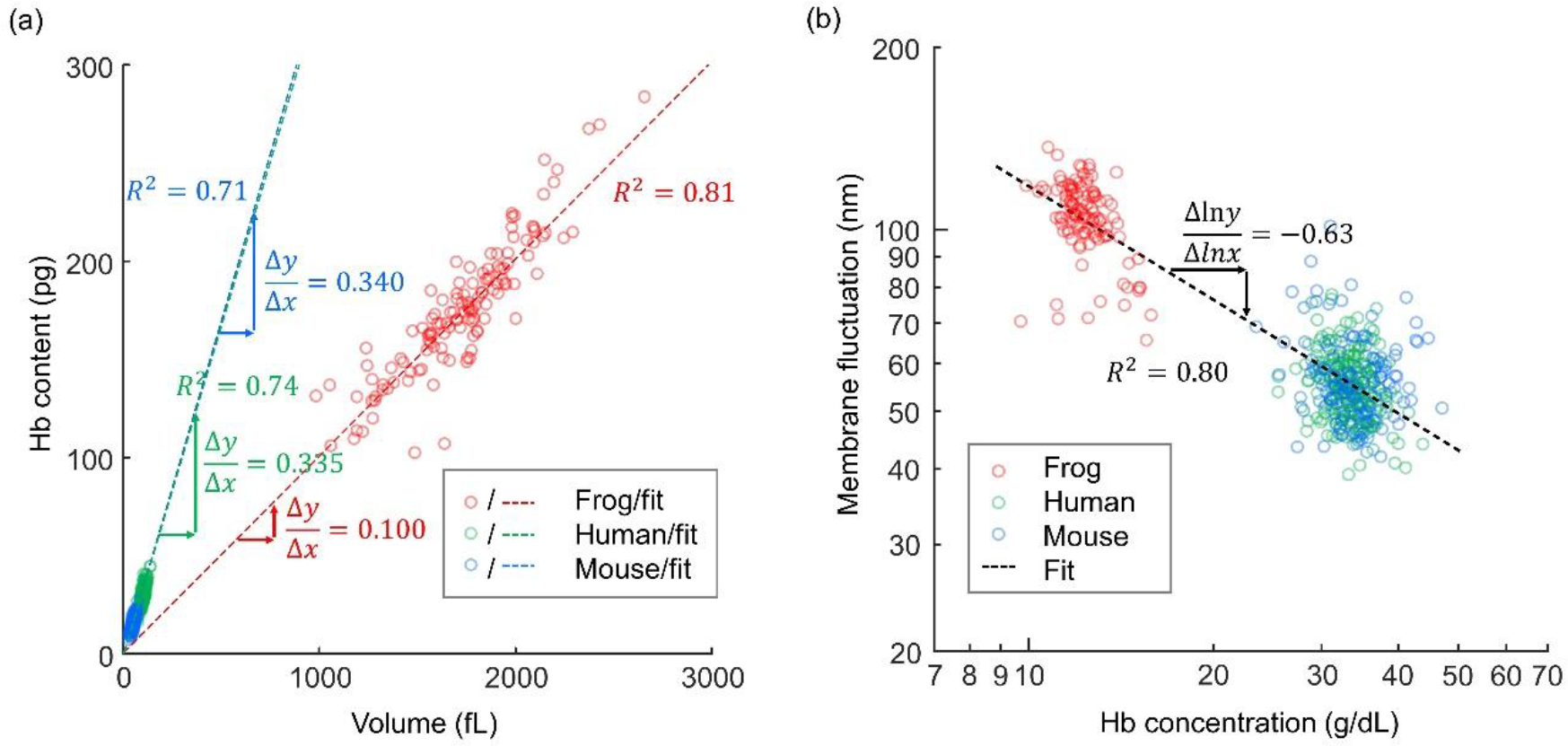
2D Scatterplots of different erythrocyte parameters. (a) A scatterplot of volumes and Hb contents. (b) A log-log scatter plot of Hb concentrations and membrane fluctuations. R^2^ indicates the R-squared value of the least square fitting.

We also performed a correlative analysis between the Hb concentration and the membrane fluctuation (Fig. 5b). In the log-log scatterplot, the membrane fluctuations of the erythrocytes from the human and mouse groups had significantly different distributions from the distributions of the frog erythrocytes. The dynamic membrane fluctuations of the frog erythrocytes (106.37 ± 14.72 nm) were significantly higher than those of the human or mouse groups (55.10 ± 6.90 nm and 57.10 ± 9.90 nm).Significantly enhanced fluctuations would be expected for the frog erythrocytes based on a simple dimensional analysis because they are 20 times greater in volume and have 3 times lower cytoplasmic viscosity. However, these large fluctuations of the frog erythrocytes are not solely explained by these two effects. For example, the presence of the nucleus and anchoring cytoskeleton may affect this mechanical property of the frog erythrocytes. The molecular organizations of erythrocyte membrane structures differ between amphibians and mammalian species. Moreover, the αI spectrin gene^50^, which is found in mammalian erythrocytes and causes enhanced deformability^51^, is absent in frog erythrocytes, and this may also explain the low deformability of the frog erythrocytes. In addition, the mechanism of dynamic remodeling of membrane cortex structures mediated by ATP^13^, which also plays a role in determining cellular deformability, may also differ in amphibian erythrocytes.

## Conclusions

In summary, we present optical measurements of 3D RI tomograms and dynamic membrane fluctuations of amphibian erythrocytes of the frog *Pelophylax nigromaculatus.* From the measured RI tomograms, morphological (cell volume and surface area), biochemical (Hb concentration and content) and biomechanical (membrane deformability) properties were quantitatively retrieved. Furthermore, a correlative imaging approach was also employed to exploit both 3D RI tomography and 3D fluorescence imaging, and to enable the investigation of nuclei structures. Using a correlative analysis, these retrieved parameters were systematically investigated and also compared with mammalian erythrocytes from humans and mice.

Although the current study focused on measurements of frog erythrocytes, the approaches used here are general and can be readily applied to the study of the erythrocytes of other species. Furthermore, the QPI methods used for the study of mammalian erythrocytes can also be applied to further understand the mechanisms of amphibian blood cells, including the label-free visualization of parasites and host cells, the visualizations of white blood cells, and the effects of ATP to cell deformability. For example, the visualization of erythrocytes in microfluidic channels *in silico* or microcapillaries of frogs and tadpoles *in vivo*^52^ would provide information valuable to unlocking the structures and dynamics of amphibian erythrocytes. In addition, this work could stimulate other disciplines including numerical simulations^53^ and analytical modeling^54^ of amphibian erythrocytes as well.

## Materials and methods

### Sample preparation

Blood samples were prepared from six female frogs *Pelophylax nigromaculatus* (*Rana nigromaculata*) which had been bred for more than 2 years in a breeding farm where the temperature and humidity were maintained over 15°C and 60% respectively. They were all fully grown, and the total length of their heads and torsos ranged from 8 to 9 cm, which is approximately the limit to which they grow. The blood was directly drawn from the heart of each frog using milliliter syringes (Sung Shim Medical, Republic of Korea) while the frogs were anesthetized with 99% ether, then immediately stored in heparin-treated vacutainers (Becton Dickinson, Franklin Lakes, U.S.A.). The extracted blood was diluted in an isotonic amphibian phosphate buffer saline solution, which was prepared by mixing 20% (w/w) deionized water and 80% (w/w) phosphate-buffered saline (PBS) solution (Gibco^®^, New York, U.S.A.).

The diluted solution was separated into unstained groups and stained groups. The unstained groups were used to collect morphological, biochemical, and mechanical erythrocyte parameters with minimal perturbation. The other stained groups were prepared by fixation with 4% paraformaldehyde solution and stained with 3 μM DAPI solution, for the purpose of cross-correlating the locations of nuclei in the erythrocytes. For imaging purposes, a 50 μl drop of erythrocyte suspension was loaded between two coverslips spaced by a strip of double-sided Scotch tape. The loaded sample was then imaged at room temperature. All the experimental protocols for the control blood donors were approved by the institutional review board of KAIST (IRB project: KH2015-37).

### Analyses of erythrocyte parameters

From the measured 3D RI tomograms of frog erythrocytes, various cellular parameters were quantitatively obtained and analyzed, including morphological (cell volume, nucleus volume, cell surface area, nucleus surface area, and sphericity index of cell shapes) and biochemical (Hb concentration and Hb content) parameters. The entire cell volume and the nucleus volume were directly retrieved using threshold values of RIs. Surface areas of the entire cells were also directly measured from the sample boundaries in 3D. To quantify the shape resemblance of a cell to a sphere, we calculated the sphericity index, which is defined as *SI* = (36*πV*^2^)^1/3^/*S*, where *V* and *S* are the entire cell volume and the surface area, respectively^55^.

The cytoplasmic Hb concentration, [Hb], was obtained using the linear relation between [Hb] and the average RI difference between the cytoplasm and the background medium, 〈Δ*n*〉, i.e. 〈Δ*n*〉 = *α*[Hb], where *α* is a RI increment^56 57^. It is known that the RI increment has a narrow range for different protein species^58^. In addition, the protein structures of Hb do not vary significantly in vertebrates^59^. Based on these two previous references, the value of a was set to have 0.18 mL/g, which was known for human erythrocytes^42,60^.

To investigate the deformability of individual frog erythrocytes, we analyzed the dynamic membrane fluctuations of the cells. The 2D holograms of erythrocytes were obtained with normal illumination at a frame rate of 125 Hz for two seconds. We assumed that the nuclear motion was stationary and its influence on the measured membrane fluctuations was negligible. From the measured optical phase delay image *Δϕ*(*x, y; t*), the height fluctuation could be calculated as *h*(*x, y; t*) = (*λ*/2*π*〈*Δn*〉)*Δϕ*(*x, y; t*). We defined the dynamic membrane fluctuation σ_*h*_ as the standard deviation of the thickness profile *h*(*x, y; t*) over time *t*.

### Statistical analysis

All the numbers that follow the ±sign in the text are standard deviations. Mann-Whitney U test using an in-built MATLAB code was used for statistical comparisons between groups.

